# From brain noise to syntactic structures: A formal proposal within the oscillatory rhythms perspective

**DOI:** 10.1101/171702

**Authors:** Mirko Grimaldi

## Abstract

The neurobiology investigation of language seems limited by the impossibility to link directly linguistic computations with neural computations. To address this issue, we need to explore the hierarchical interconnections between the investigated fields trying to develop an inter-field theory. Considerable research has realized that event-related fluctuations in rhythmic, oscillatory EEG/MEG activity may provide a new window on the dynamics of functional neuronal networks involved in cognitive processing. Accordingly, this paper aims to outline a formal proposal on neuronal computation and representation of syntactic structures within the oscillatory neuronal dynamics. I briefly present the nature of event-related oscillations and how they work on the base of synchronization and de-synchronization processes. Then, I discuss some theoretical premises assuming that reentrant (hierarchical) properties of synchronized oscillatory rhythms constitute the biological endowment that allow the development of language in humans when exposed to appropriate inputs. The main rhythms involved in language and speech processing are examined: i.e. theta, alpha, beta, and gamma bands. A possible formal representation of the syntactic structures on the base of these oscillatory rhythms is discussed: in this model, the theta-gamma rhythms are cross-frequency coupled into the alpha-gamma-beta and into the gamma-beta-theta rhythms to generate the sentence along reentrant cortico-thalamic pathways through Merge, Label and Move operations. Finally, I present few conclusive remarks within an evolutionary perspective.

> *“…The whole burden of philosophy seems to consist in this, from the phenomena of motions to investigate the forces of nature, and from these forces to demonstrate the other phenomena.”* (Newton, 1687/1726)

## 1 Introduction

The neurobiology investigation of language seeks to uncover the relation between the linguistic computations and its representations in the brain. In doing this, we need to coherently correlate linguistic ontologies – e.g., phoneme, syllable, morpheme, lexicon, syntax and their operations – with neurophysiological ontologies – e.g., neuron, dendrites, spines, synapses, action potentials and their operations. This is a not effortless task, since the two entities seem not directly commensurable. Furthermore, linguistic computation involves a number of fine-grained levels and explicit computational operations – that is how phonemes are combined together to form syllables and words, and how words are combined together to form sentences – whereas neuroscientific approaches to language operate in terms of broader conceptual distinctions (e.g., what areas of brain are deputed to phonology and what to syntax, etc.). These represent what Poeppel & Embick (2005) call, respectively, the Ontological Incommensurability Problem and the Granularity Mismatch Problem (see also Embick & Poeppel 2015; Grimaldi 2012). Thus, a direct reduction of the linguistic primitives into neurobiological primitives is a limitation in the progress of an integrated study of language and brain.

These issues may be solved if we assume that our description of the world is founded on various hierarchies. At one end we have concepts and words we use to capture some facts of the world, at the other end we have the fundamental laws of physics (Feynman 1967). For example, when we say ‘heat’, we are using a word for a mass of atoms which are jiggling, and when we say ‘salt of crystal’ fundamentally we are referring to a lot of protons, neutrons, and electrons. So, we may describe the world using ordinary language ignoring the fundamental laws. When we go higher up from this, we found words as ‘phoneme’ or ‘syntax’ to capture some computational properties of human language: (i) the fact that we use contrastively specific acoustic-articulatory features of sounds to generate words (as, for instance, the sounds [k] and [r] in [‘kæt] *cat* vs. [‘ræt] *rat*); (ii) and the fact that sentences are characterized by particular relation among words, also at long distance: e.g., *The book that was lying under all the other books is the most interesting*. As we go up in this hierarchy of complexity, we get words as ‘neurons’ and ‘synapse’ that refer to sophisticated chemical and electrical processes in the physical world and that control such computational properties of human language. In brief, we use different concepts and notions (or ontologies) to understand the world at an ever higher level.

How to correlate these different levels of the world knowledge? The best way is to investigate the world at various hierarchies, looking at the whole structural interconnection of the levels. So, we cannot draw carefully a line all the way from one end of the hierarchy to the other looking at the world in term of monolithic entities “[…] because we have only just begun to see that there is this relative hierarchy […]. The great mass of workers in between, connecting one step to another, are improving all the time our understanding of the world, both from working at the ends and working in the middle, and in that way we are gradually understanding this tremendous world of interconnecting hierarchies” (Feynman 1967: 125-126).

I think this view presupposes the development of an inter-field theory that integrates and bridges fields rather than establishing one complete, unified theory. Inter-field theories can be generated when two fields share an interest in explaining different aspects of the same phenomenon in order to build solid knowledge and relations between the fields. This perspective advocates integration rather than reduction (see for example Murphy & Benitez-Burraco 2017). Accordingly, an inter-field theory should interconnect well-established linguistic computational primitives with neurophsysiological computations responsible for representational processes at the light of the knowledge reached within each research area. This demanding task will lead us to progressively create epistemological bridges between different disciplines. More precisely, the task ahead is to characterize this kind of linked computations and find out how they work in concert producing linguistic behaviors: and step by step it is probable that the ‘neurobiology of language’ may stand on its own feet integrating the two research traditions and producing an inter-theoretic framework.

Recently, considerable research has achieved that event-related fluctuations in rhythmic, oscillatory electroencephalography (EEG)/magnetoencephalography (MEG) activity may provide a new window on the dynamics of the coupling and uncoupling of functional neuronal networks involved in cognitive processing (Sauseng & Klimesch 2008; Canolty et al. 2010; Donner & Marcus 2011; Hanslmayr et al. 2016). This perspective, as we will see, offers the possibility to directly explore the interconnection between linguistic ontologies and neuronal ontologies. In this work, I aim to sketch a proposal on neuronal computation and representation of syntactic structures within the oscillatory neuronal dynamics along the line of previous studies (Murphy 2015a, 2016; Boeckx & Theofanopoulou 2014). Therefore, I will present and discuss a new representation of syntactic structures reinterpreting the classical tree-diagram according to oscillatory rhythms principles.

## 2 Electroencephalography, event-related potentials, and event-related oscillations

Electrodes placed in different areas of the scalp provide recording of the brain’s electrical activity and noninvasive sensitive measures of brain functions in humans. Regardless of whether an individual receives sensory information or performs higher cognitive processes, the brain exhibit measurable electrical activity. By recording this activity with numerous electrodes, researchers have developed different approaches to determine when (and at least where) in the brain information processing occurs.

A first approach uses to monitor neural phenomena in the continuous EEG/MEG recording of brain activity when the subject is at rest and not involved in a task. It reveals the sum of the random activity of millions of neurons that have similar spatial orientation in the brain. This activity typically fluctuates in wave-like patterns, and depending on the frequency of these patterns, one distinguishes different brain waves called: delta (~0.5-4 Hz), theta (~4-10 Hz), alpha (~8-12 Hz), beta (~12-30 Hz), and gamma (~30-100 Hz) rhythms. Traditionally, variations in the patterns of these brain waves can indicate the level of consciousness, psychological state, or presence of neurological disorders.

A second approach consists to record the EEG/MEG while subjects are performing a sensory or cognitive task. Thereby stimuli are presented to subjects and markers are set into the EEG trace whenever a stimulus is presented. Then a short epoch of EEG/MEG around each marker is used to average all these segments. This is based on the logic that in each trial there is a systematic brain response to a stimulus. Practically, this means that one typically repeats a given experimental paradigm a number of times (say, >30 times), and then one averages the EEG/MEG recordings that are recorded time-locked to the experimental event. However, this systematic response cannot be seen in the raw EEG, as there it is overlaid by a lot of unsystematic background activity (which is simply considered as noise). By averaging all the single epochs that are time-locked to the experimental event, only the systematic brain response should remain (i.e., those generate neural action potentials related to the stimuli), but the background EEG/MEG should approach zero (Sauseng & Klimesch 2008). The noise (which is assumed to be randomly distributed across trials) diminishes each time a trial is added to the average, while the signal (which is assumed to be stationary across trials), gradually emerges out of the noise as more trials are added to the average. These brain responses are named event-related potentials (ERPs) and event-related magnetic fields (ERMFs) reflecting the summated activity of network ensembles active during the task.

ERPs/ERMFs are characterized by specific patterns called ‘waveforms’ (or ‘components’), which are elicited around 50-1000 ms starting from the onset of the stimulus and show positive (P) and negative (N) oscillatory amplitudes (i.e., voltage deflections). For instance, P100, N100, P200, P300, N400, P600 are the principal components elicited during language processing starting from sound perception to semantic and syntactic operations. So, this technique provides millisecond-by-millisecond indices of brain functions and therefore provide excellent temporal resolution.

It is important to realize that the amplitude of an oscillation is, roughly speaking, the size of its (positive or negative) peak deflection relative to some baseline (that is, how big the oscillation is). There is, however, another notion that we need to consider: the phase of an oscillation. Roughly speaking, the phase is the slope (or direction) of the signal at a given one point in time, which is equivalent to the left–right shift of the oscillation. In this respect, ERPs are time- and phase-locked to the event (i.e., the experimental stimuli) that generated the oscillatory activity.

Although the ERP approach has opened an important window on the time course and the neural basis of speech and language processing, more than 100 years after the initial discovery of EEG activity, researchers are turning back to reconsider another aspect of EEG, that is the event-related oscillations. This is because an increasing number of researchers began to realize that an ERP only represents a certain part of the event-related EEG signal. Actually, there is another aspect of extreme interest for the study of cognitive functions: the event-related fluctuations in rhythmic, oscillatory EEG/MEG activity. This view, indeed, might provide a new window on the dynamics of the coupling and uncoupling of functional networks involved in cognitive processing (Varela et al., 2001). In fact, substantial literature now indicates that some ERP features may arise from changes in the dynamics of ongoing EEG rhythms/oscillations of different frequency bands that reflect ongoing sensory and/or cognitive processes (Başar 1999; Başar et al. 2001; Buzsaki 2006). More precisely, the EEG oscillations that are measured in a resting state become organized, amplified, and/or coupled during cognitive processes. It has been argued that ERP does not simply emerge from evoked, latency–fixed polarity responses that are additive to and independent of ongoing EEG (Sauseng et al., 2007): instead, evidence suggests that early ERP components are generated by a superposition of ongoing EEG oscillations that reset their phases in response to sensory input, (i.e., the external or internal stimuli generating cognitive activities). Therefore, event-related oscillation, further than to have the time-locked EEG information, permits to retrieve the non-phase locked EEG information related to the cognitive activity induced by the stimulus.

Within this perspective, ongoing cerebral activity can no longer be thought of as just relatively random background noise (the non-phase EEG activity) that must be removed in order to see the event-related responses, but as a whole containing crucial information on the dynamical activity of neural networks: thus, the EEG and ERP are the same neuronal event, as the ERP is generated because of stimulus-evoked phase perturbations in the ongoing EEG. A fundamental feature of the phase-resetting hypothesis is that following the presentation of a stimulus, the phases of ongoing EEG rhythms are shifted to lock to the stimulus. From this, it follows that during pre-stimulus intervals, the distribution of the phase at each EEG frequency would be random, whereas upon stimulus presentation, the phases would be set (or reset) to specific values (for each frequency). The resetting of the phases causes an ERP waveform to appear in the average in the form of an event-related oscillation (Makeig et al. 2002; Penny et al. 2002; Klimesch et al. 2004).

Unlike ERP (based on the analysis of components), event-related oscillation is based on the time-frequency analyses (e.g., Gross, 2014). One such method is wavelet analysis.

The general idea is that not all relevant EEG activity is strictly phase-locked (or evoked) to the event of interest (Buszáki, 2006). Obviously, this activity shortly before stimulus onset is mostly not visible in ERPs due to cancellation; nevertheless, this pre-stimulus baseline activity may have a crucial impact on the observed ERPs (Klimesch, 2011). Time-frequency analyses enable us to determine the presence of oscillatory patterns in different frequency bands over time. Thus, with wavelet analyses, it can be established whether oscillatory activity in a specific frequency band, often expressed in power (squared amplitude), increases or decreases relative to a certain event, as represented in Fig. 1.

**Fig. 1:**
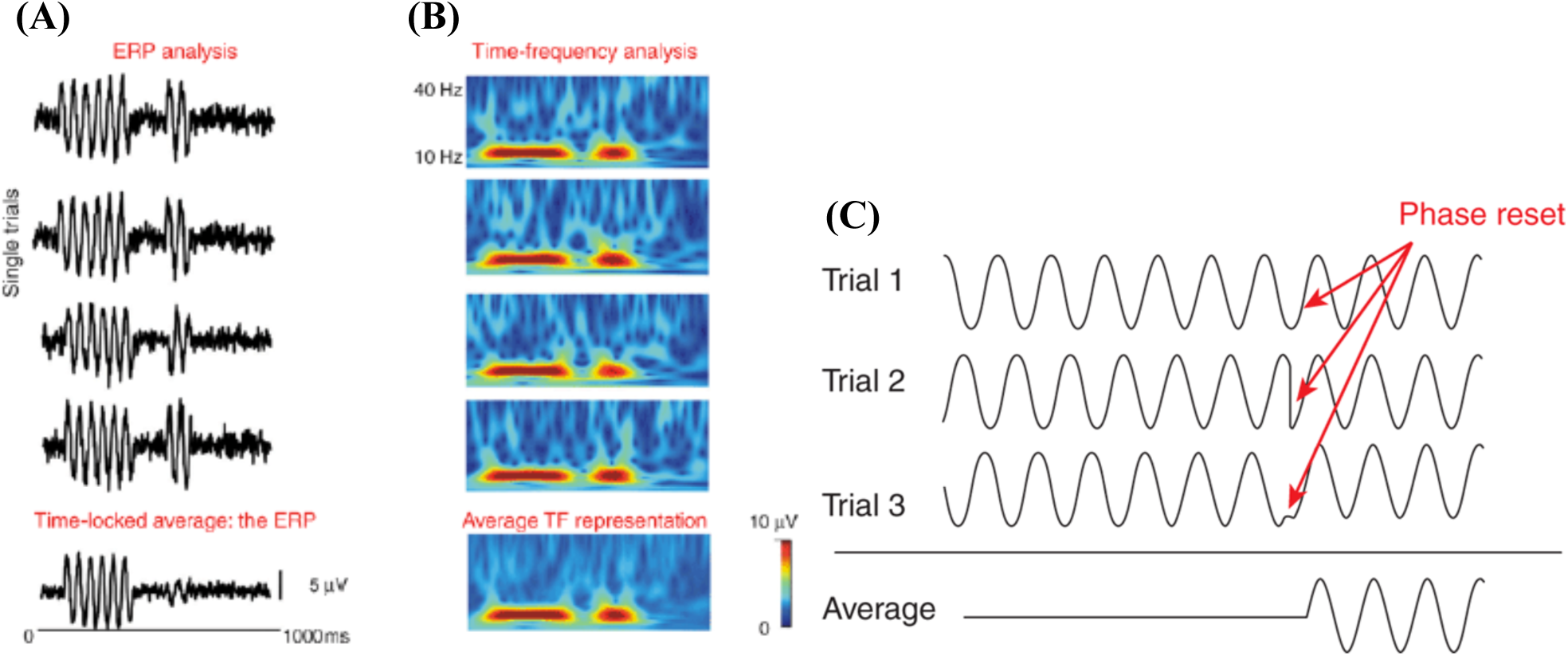
Simulated EEG data illustrating the difference between phase-locked (evoked) activity and non-phase-locked (induced) activity. (A): Single-trial EEG time courses showing two consecutive event-related responses (an amplitude increase at 10 Hz). The first response is phase-locked with respect to the reference time-point (t=0), and as a result this evoked response is adequately represented in the average ERP. The second response is time-locked, but not phase-locked to t=0, and as a result this induced response is largely lost in the average ERP. (B): time-frequency (TF) representations of each single trial, with red colors coding for the amplitude increase at 10 Hz. Crucially, the average TF representation contains both the phase-locked and the non-phase-locked responses. (C): simulated data illustrating the principle of phase resetting. Three single trials are shown whose phases are not aligned initially. Red arrows indicate the point in time at which an event-induced phase reset occurs. The bottom trace shows what the average ERP would look like if a sufficient number of such trials (in practice >30 trials) are averaged. Adapted from Bastiaansen, Mazaheri & Jensen (2012).

The importance in considering the non-phase locked event-related oscillations consists in the fact that, contrary to phase-locked responses as ERPs, they reflect the extent to which the underlying neuronal activity synchronizes. Synchronization and de-synchronization are related to the coupling and uncoupling of functional networks in cortical and subcortical areas of the brain (see, e.g., Varela et al., 2001). This aspect, of course, is related to how different types of information, which are stored in different parts of the network, are integrated during computational and representational processes. Importantly, elements pertaining to one and the same functional network are identifiable as such by the fact that they fire synchronously at a given frequency. This frequency specificity allows the same neuron (or neuronal pool) to participate at different times in different representations. Hence, synchronous oscillations in a wide range of frequencies are considered to play a crucial role in linking areas that are part of the same functional network. Importantly, in addition to recruiting all the relevant network elements, oscillatory neuronal synchrony serves to bind together the information represented in the different elements (Gray et al. 1989).

## 3 Theoretical premises

Inspired by Başar (2011), I assume that cognitive computational and representational processes are intrinsic to brain oscillatory activity. This oscillatory activity is characterized by coherent cooperation between distant structures through different oscillatory phases. Thus, according to Lasheley (1929), the brain operates as a “whole” thanks to rhythmic oscillations that are selectively distributed in the whole brain: it is the coordination and coherence of oscillations that generate parallel sensory-cognitive processing. Research has shown that neural population in cortical and subcortical areas (e.g., cortex, hippocampus or cerebellar cortex) are all tuned to the very same frequency ranges (Steriade et al., 1992; Başar, 1999). These findings support the hypothesis that all brain networks communicate by means of the same set of frequency codes of rhythmic oscillations. This presupposes that the intrinsic oscillatory activity of each single neuron shapes the natural frequencies of neural assemblies, that is the delta, theta, alpha, beta, and gamma frequencies.

As noted above, ERP components seem generated by a superposition of ongoing EEG oscillations that reset their phases in response to sensory input: this superposition principle suggests that there exists synergy between oscillations during performance of sensory-cognitive tasks. Accordingly, integrative brain function necessary for sensory-cognitive processing may be obtained through the combined action of multiple oscillations. Also, the superposition principle is crucial for memory functions directly correlated with all brain functions, and, in particular, with speech and language functions. This is a crucial point, because memory-related oscillations must have dynamic properties evolving in different hierarchical states that take place along a continuum where the boundaries of memory states integrate into each other. In line with Murphy (2015a), I assume that such property plays a key role in the basic computations that characterize the Faculty of Language: that is, Merge, Label, Move and its correlated Spell-Out operations (Chomsky 1995, 2001, 2013). Actually, these computational operations need that ‘mnemonic objects’ – concepts, words and related information determining, for example, agreement and case, etc. – are dynamically manipulated thanks to bidirectional exchange of signals along reciprocal axonal fibers linking two or more brain areas from thalamus to cerebral cortex and back.

According to Edelman (1989, 1993, 2004) a large and diverse body of evidence suggests that intermittent signaling along reentrant paths is critical to a variety of neural functions in vertebrate brains, ranging from perceptual categorization to motor coordination and cognition. Reentry takes on a variety of forms enabling many different processes. These processes facilitate the coordination of neuronal firing in anatomically and functionally segregated cortical areas. By these means they bind cross-modal sensory features by synchronizing and integrating patterns of neural activity in different brain regions. Reentrant signaling is a ubiquitous and dominant structural and functional motif of vertebrate telencephalons. Reentry has, conversely, rarely, if ever, been characterized in an invertebrate nervous system, and it may be a relatively recent evolutionary innovation. Reentrant processes are those that involve one localized population of excitatory neurons simultaneously both stimulating, and being stimulated by, another such population: the structural architecture that generates this process is likewise referred to as reentrant. Experimental evidence converges to indicates that processes of reentry play widespread and essential roles in vertebrate brain function, evolution, and development (Edelman & Gally 2013). The reciprocal exchange of signals among neural networks in distributed cortical and cortico-thalamic areas – when combined with appropriate mechanisms for synaptic plasticity – results in the spatiotemporal integration of patterns of neural network activity (cf. Fig. 2). This process may be considered a kind of neural recursion that allows the brain to categorize sensory input, remember and manipulate mental objects, generate motor commands and/or cognitive activity. In particular, it has been suggested that the hippocampal declarative memory system may control cognitive functions that require on-line integration of multiple sources of information, such as on-line speech and language perception and production processing (Duff & Brown-Schmidt 2012).

**Fig. 2:**
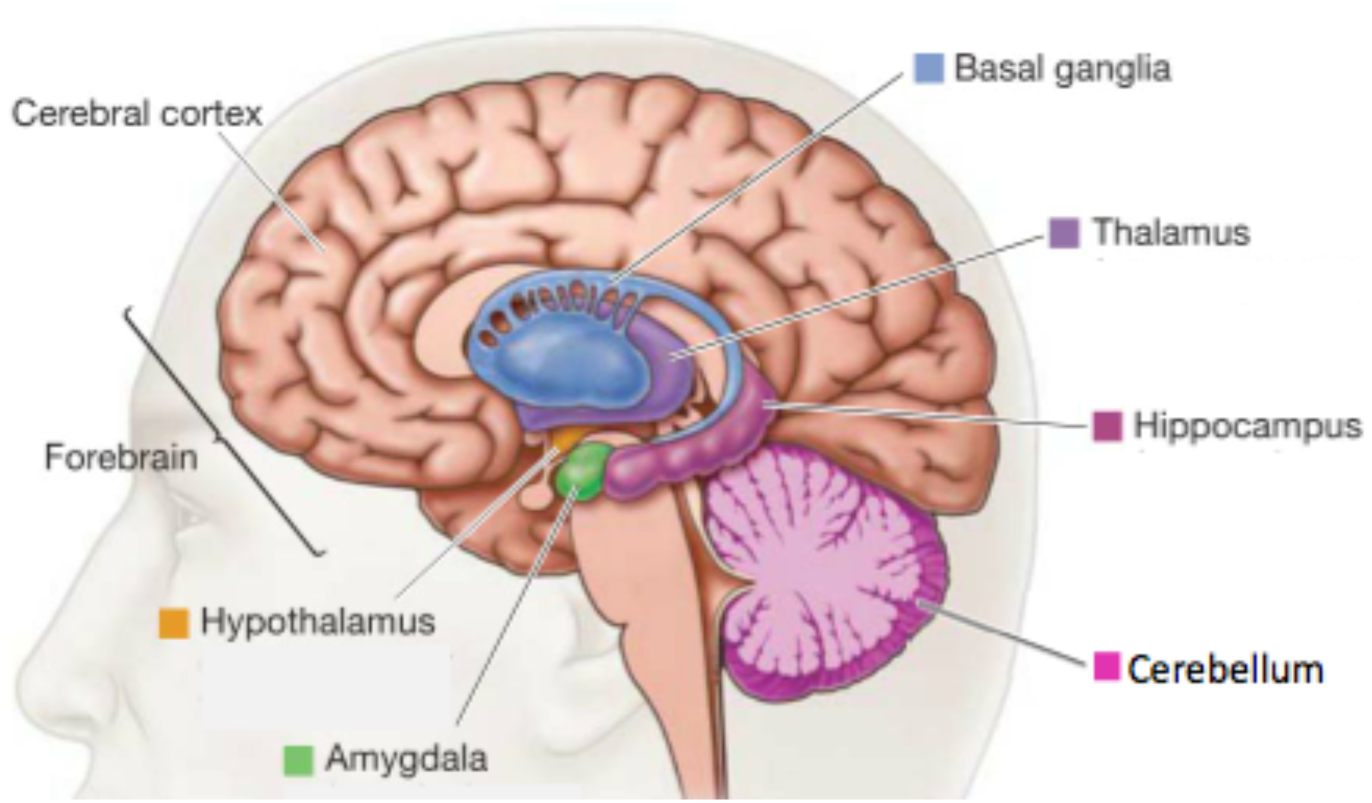
Cortical and cortico-thalamic structures forming reentrant synchronized pathways. Adapted from http://www.proprofs.com/flashcards/story.php?title=intro-mind-and-brain‐‐topic-2-foundation-brains.

Within the cortico-thalamic pathway, the basal ganglia assume strategic function for speech and language processing. Thanks to the thalamus, the basal ganglia, the cerebellum, and the hippocampus interface with the cortex in a reciprocal fashion (Theofanopoulou & Boeckx 2016; Hickok 2012). The basal ganglia process information indirectly in a set of loops, whereby they receive input from the cortex and return it to the cortex via the thalamus. In that way, the basal ganglia modify the timing and amount of activity that leaves the cortex and travels down the pyramidal pathway effectively modulating the neural activity for motor and cognitive processes (cf. Fig. 2).

In particular, basal ganglia dysfunction in humans can result in a subcortical dementia where an afflicted individual will perseverate, finding it difficult, in some cases impossible, to change the direction of a thought process (Flowers & Robertson 1985), or comprehend sentences that has moderately complex syntax (Lieberman et al. 1992). Furthermore, the basal ganglia are also involved in associative learning (Lieberman 2009). Although it is clear that basal ganglia are not directly involved in core semantic operations (their lesion does not generate semantic syndrome), they are recruited in the intention to retrieve lexical items during word generation regardless of the semantic category (Crosson, Benjamin, Levy 2007).

In generating sentences, we access to the knowledge system (KS): that is, the complex of processes represented by the long-term memory system together with the procedural and perceptual system (Klimesh 2012). The basic idea is that the KS interacts with the working memory (a multi-component system that holds and manipulates information in short-term memory) in a way that traces stored in the KS are used for short-term storage (Klimesch, Schack 2003). This is possible thanks to the synchronization property of neurons according to which different brain (cortical and sub-cortical) regions may be synchronized through phase amplitude cross-frequency coupling whereby phases of lower frequencies modulates the power of higher frequencies: for instance, the coupling between the phase of theta and the power of gamma (Hanslmayr et al. 2016).

I assume that synchronization reflects a basic computational principle that underlies the dynamic control of effective interactions along selective subsets of the anatomically possible neuronal connections. In other words, selectively distributed oscillatory rhythms act as resonant communication networks through large populations of neurons, with functional relations to memory and integrative functions. The implication is that the cross-frequency synchronization between oscillatory rhythms reflects the interaction between working memory and the KS. Thus, the access to the KS and the computations generated are a continuous process dynamically organized; furthermore, the memory functions from the simplest sensory memories to the most complex semantic and episodic memories are manifested in distributed multiple oscillations in the whole brain. On this biological mechanism is grounded the acquisition of natural languages. The reentrant (hierarchical) properties of synchronized oscillatory rhythms constitute the biological endowment that allow the development of grammar in human beings when exposed to some appropriate inputs. Inputs generating computations and representations are structured in memory according to universal biological constrains and some degrees of freedom (options) that the neural system presents: these optionality is at the basis of variation (and micro-variation) characterizing natural languages.

## 4 From oscillatory rhythms to syntactic structures

### 4.1 Functionality of rhythms for language and speech processing

Theta oscillations can be found in the human cortex, the hippocampus, and the hypothalamus. Theta oscillations seem to be important for a variety of cognitive functions. It was shown that hippocampal and cortical theta activity is associated with virtual navigation, declarative memory processes, successful memory encoding, the amount of information held in memory, and episodic memory processing (Sauseng & Klimesch 2008). Theta power increases during language processing have been related to the retrieval and encoding of lexical semantic information (Bastiaansen, et al. 2008; Bastiaansen & Hagoort 2015). Additionally, working memory-load-dependent increase of theta activity has been suggested: i.e., when the amount of encoded information increases, theta activity grows stronger (Jensen & Tesche 2002). Weiss et al. (2005) found higher anterior–posterior theta coherence over the left hemisphere during the processing of relative clauses and suggest it may be related to the initiation of linguistic analysis since coherence during linguistic analysis is higher in the left hemisphere. Thus, it seems that theta activity could be a correlate of control processes when multiple items have to be held in working memory to be managed or bound in comprehensive memory entry (Lisman & Idiart 1995). Indeed, recent studies suggest that higher gamma frequency oscillations can be nested into theta cycles. This seems to reflect organization of multiple items into sequential working memory representations or integration between sensory bottom-up and top-down memory representations (Sauseng et al. 2010).

Gamma oscillations, on the other hand, are cortically generated and arise from intrinsic membrane properties of interneurons or from neocortical excitatory-inhibitory circuits (Sauseng & Klimesch 2008). Actually, synchronization phenomena of this brain rhythm were related to binding of information. More recently, effects at human gamma frequency were also reported for the encoding, retention and retrieval of information independent of sensory modality. It has also been discussed that gamma binds large-scale brain networks (Kahana, 2006). Recently, Bastiaansen & Hagoort (2015) have clearly showed that gamma band neuronal synchronization is involved in sentence level semantic unification operations. This gamma-band effects have maxima over the left posterior temporal and the left frontal scalp, which is well compatible with the notion that semantic unification is a result of a dynamic interplay between left posterior superior/medial temporal gyrus and inferior frontal gyrus. Interestingly, a recent ECoG study (Rapela 2016) showed that rhythmic speech production (i.e., sequence of syllables) modulates the power of high-gamma oscillations over the ventral sensory motor cortex, a cortical region that controls the vocal articulators, and the power of beta oscillations over the auditory cortex (due to the auditory feedback necessary control acoustic-articulatory outputs). He found significant coupling between the phase of brain oscillations at the frequency of speech production and their amplitude in the high-gamma range (i.e., phase-amplitude coupling, PAC). Furthermore, the data showed that brain oscillations at the frequency of speech production were organized as traveling waves and synchronized to the rhythm of speech production.

The functional relevance of alpha oscillations is very widespread. There is strong evidence that alpha amplitudes are related to the level of cortical activation. A strong alpha activity is associated with cortical deactivation or inhibition, but it is also involved in highly specific perceptual, attentional, and executive processes functions in working memory processes as in responding selectively to semantic task demands (Klimesch et al. 2005; Bartsch et al. 2015). Actually, Klimesch (2012) argues that alpha-band oscillations reflect the temporal structure of one of the most basic cognitive processes, which may be described as ‘knowledge-based consciousness’ and which enables ‘semantic orientation’ via controlled access to information stored in the knowledge system. Furthermore, Benedek et al. (2011) found frontal alpha synchronization during convergent and divergent thinking only, under exclusive top-down control (high internal processing demands), suggesting that these rhythms are related to high internal processing demands which are typically involved in creative thinking. Finally, Strauβ et al. (2015) demonstrated that alpha phase – both before and during the presentation of word or word-like stimuli – predicts the accuracy of lexical decisions in noise.

As alpha rhythms, also beta oscillations are cortically generated, due to their local strictness, although widespread cortical beta networks in humans have been shown (Gross et al. 2004). From a functional perspective beta oscillations have mainly been associated with motor activity, but beta has also been suggested to play an important role during attention or higher cognitive functions (Razumnikova 2004), as for binding mechanisms during language processing (Weiss & Mueller 2012). Indeed, it was also proposed that beta frequencies are used for higher-level interaction between multimodal areas involving more distant structures and the binding of temporally segregated events, which is especially important for language processing (Donner & Siegel 2011; Lam et al. 2016). Generally, it has been shown that target words for syntactically (Davidson & Indefrey 2007; Bastiaansen et al. 2010; Pérez et al. 2012; Kielar et al. 2014) and semantically (Luo et al. 2010; Wang et al. 2012; Kielar et al. 2014) acceptable sentences beta power was higher than target words resulted in syntactic or semantic incongruities. Accordingly, Lewis & Bastiaansen (2015) and Lewis et al. (2016) suggest that the increased beta activity reflects the active maintenance of the current neurocognitive network responsible for the construction and representation of the sentence-level meaning. It may also indicate a greater reliance on top-down predictions based on that sentence-level meaning (i.e., the increased activity may be related to greater weighting of the top-down signal based on the current generative model), in order to actively try to integrate the new linguistic input into the current sentence-level meaning representation. Along this line, beta synchronization has been correlated with the binding of semantic features of different lexical categories (Weiss & Mueller 2003). Crucially, Bastiaansen & Hagoort (2015) performed an elegant experiment showing that beta-band power is strictly related to syntactic structure building at the sentence level with a maximum around the vertex. Weiss et al. (2005), on the other hand, suggest that while theta changes may be associated with memory processes and gamma with attentional effort, beta bands may be activated with semantic–pragmatic integration.

All in all, (i) theta rhythms seem involved in retrieving lexical semantic information and controlling processes with multiple items. This process may be supported by the nesting of gamma frequency oscillations into theta cycles reflecting organization of multiple items into sequential working memory representations or integration; (ii) gamma-band neuronal synchronization on the one hand seems related to sentence level semantic unification, on the other hand to speech production; (iii) alpha phase acts not only in decisional weighting, but also in semantic orientation, in creative thinking, lexical decisions; (iv) beta synchronization serves to bind distributed sets of neurons into a coherent representation of (memorized) contents during language processing, and, in particular, to building syntactic structures. It is important to note that it is impossible to assign a single function to a given type of oscillatory activity (Başar et al. 2001). It is thus unlikely that, for instance, theta has a single role in language processing. In fact, theta’s role and its varying patterns of coherence as a function of task demands may be better seen in its relationship to beta and gamma (and same thing is true for the other oscillatory rhythms). Accordingly, it may be important to consider the simultaneous changes in the coherence patterns in the different frequency ranges.

### 4.2 An inter-field model for syntactic structure generation

The challenge now is to develop an inter-field model that coherently interconnect and integrate neural computations (i.e., those intrinsic to oscillatory rhythms) with syntactic computations assumed as primitives within linguistic theory. The best candidate to sketch an interconnected neurobiological model is the fundamental structure-building operation of natural language syntax (Chomsky 1995, 2002, 2013): i.e., Merge and Label – or, according to Hornstein (2009) – Concatenation plus Label. Merge is an operation that takes a number of syntactic objects (lexical items) and join them together to form a unit. Merge strings together two elements when one selects the other and the element which projects (assigns the label to the whole structure) is the selector. The selector is also called the “head” of the construction. Note that Merge is recursive: so, we can string together multiple instances of Merge to create ever larger structures. For instance, image to realize the sentence in (1):

#### (1) The cat lays on the carpet

Merge takes the two lexical items *the* and *cat* and form a new object: [the cat]. The new unit has a label, which is inherited from one of the merged elements, i.e. the determiner, forming the Determiner Phrase (DP) [_DP_ the cat]. The lexical items *on the carpet* fall on the same operation: in this case the element that assigns the label to the structure is the preposition *on* forming the Prepositional Phrase (PP) [_PP_ on the carpet] which also contains the DP [_DP_ the carpet]. The item *lays* is then merged with the PP [on the carpet] generating the Verbal Phrase (VP) [_VP_ lays on the carpet]. The VP is merged with the DP [_DP_ the cat]. Then the VP is merged with the Inflectional Phrase (IP) – a phrase that have inflectional properties – generating the sentence in (1): [_IP_ [_DP_ the cat [_VP_ lays [_PP_ on [_DP_ the carpet]]]]].

According to the picture above outlined, I assume that cyclic, dynamic, and hierarchical oscillations cross-frequency coupling synchronize sub-cortical and cortical regions generating a functional neuronal network and ensuring bottom-up and top-down computations and communication. The strength of phase-amplitude cross-frequency coupling differs across brain areas in relation to cognitive processes accomplished: while high-frequency (beta, gamma) brain activity reflects local domains of cortical processing, low-frequency (theta, alpha) brain rhythms are dynamically entrained across distributed brain regions by both external sensory input and internal cognitive events. Thus, cross-frequency coupling may serve as a mechanism to transfer information from large-scale brain networks operating at behavioral timescales to the fast, local cortical processing required for effective computation and synaptic modification, integrating functional systems across multiple spatiotemporal scales (Canolty & Knight 2010).

In generating a sentence, I suggest the following neuronal operations controlled by oscillatory rhythms in generating syntactic structures:

– First of all, a speaker needs to access to the knowledge system where long-term memory interacts with working memory in retrieving conceptual objects (within the hippocampal and hypothalamic structures) mapping them onto lexical items (at the cortical level within fronto-temporal structures): this step is controlled by nested *theta*-*gamma* oscillations which organize multiple items into sequential representations between sensory bottom-up and top-down memory representations. So, following Jensen and Lisman (1998) and Murphy (2016a), I postulate that theta and gamma interact in the process of storing lexical representations in declarative memory.
– Cyclic coupling of *alpha*-*gamma*-*beta* rhythms are involved in merging and labelling lexical items. In particular, alpha oscillations (implicated in lexical decision) control what lexical items, selected to realize an appropriate sentence, may be grouped into units identifying phrases typologies; gamma rhythms, on the other hand, rule the overall process of merging and labelling (this process probably needs high frequency oscillation in order to rapidly control and concatenate a large number of lexical items, where also morphological information are computed); finally, beta bands control processes concerning inflectional properties of the syntactic structures generating a coherent representation of sentences. These rhythms are likely responsible for the Spell-Out transfer operation to conceptual-intentional (CI) interface.
– *Gamma*-*beta*-*theta* oscillations are deputed to supervise the Spell-Out transfer to sensory-motor (SM) interface. More precisely, this cross-frequency bands may control the production of sequence of syllables: they may have a crucial role in cyclic interacting with long-term memory and working memory. Actually, for what concerns speech perception processing, it has been suggested that a remarkable correspondence between average durations of speech units and the frequency ranges of cortical oscillations exists: phonetic features are associated with high gamma and beta oscillations, and syllables and words with theta oscillations (Giraud & Poeppel 2012). So, I hypothesize that the same mechanism can be reflected at the production level to control the speech perception-production interface (as recent data suggest: Rapela 2016). In other words, the conceptual objects retrieved by speakers from long-term memory – where I hypothesized the theta-gamma oscillation are involved – should contain not only lexical and morpho-syntactic information but also phonetic and phonological information. At a certain point of the computational and representational processes the former are Spelled-Out to CI interface and the latter to SM interface resorting to cyclic cross-frequency synchronization.

For what concerns the SM interface, it should be interesting for future research to test whether also delta bands are involved a suggested by recent perceptive data (Giraud & Poeppel 2012). In fact, low-frequency oscillations at the delta (1–3 Hz) band seem to correspond to slower modulations such as phrase level prosody (Ghitza 2011). It is well known, indeed, that prosodic patterns drive the syntactic derivation and the formation of the prosodic representation in compliance with the T-model of grammar (e.g., see Bocci 2013; Hauser, Chomsky, Fitch 2002).

The formal proposal here outlined may be represented through a neuronal tree, as illustrated in Fig. 3. The hypothesis is that the neuronal tree has a bottom-up generation in line with the idea that sub-cortical oscillations are cyclically and selectively cross-frequency structured with upper cortical oscillations forming a neuronal network and allowing the brain to operate as a “whole” in real time: as above noted, this mechanism ensures bottom-up and top-down computations. If we want to develop an inter-field model aiming to integrate neural computation with those computations assumed to play a crucial role in syntactic structures, we need to link the neuronal primitives with the linguistic primitives as much as possible.

**Fig. 3:**
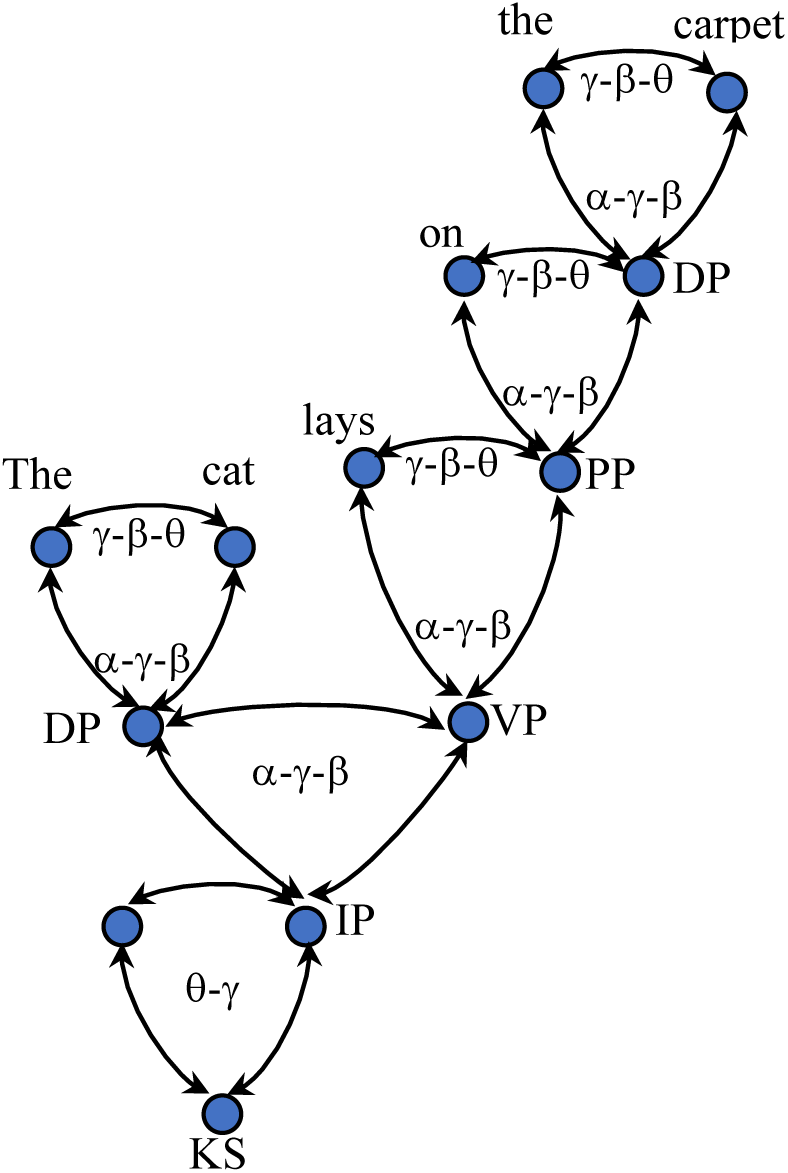
Neuronal tree representation of the sentence *The cat lays on the carpet*. Hypothesized oscillatory rhythms involved in computational and representational processes are highlighted. KS = knowledge system; IP = Inflectional Phrase; DP = Determiner Phrase; VP = Verbal Phrase; PP = Prepositional Phrase. Starting from the KS, the θ-γ rhythms are cross-frequency coupled into the α-γ-β and into γ-β-θ rhythms to generate the sentence along reentrant cortico-thalamic pathways through Merge, Label and Move operations. The vertical arrows between the nodes indicate the ascending information conveyed by thalamic nuclei while the horizontal arrows between the nodes indicate the descending information from fronto-temporal cortical areas.

Accordingly, I propose to start representing the retrieval of the conceptual objects from the KS (in the thalamic nuclei) where cross-frequency coupling of theta-gamma oscillations evaluate and broadly organize them into lexical times. Then, phase resetting (cf. Fig. 2) synchronizes the coupling of alpha-gamma-beta rhythms generating Merge, Label and Inflectional computations together with the representation of the structure (a transfer to CI interface is possible at this point). Finally, a subsequent phase resetting into gamma-beta-theta rhythms projects syntactic structures to SM interface. The arrows in Fig. 1 represent the idea that computational and representational processes are guided by a cyclic principle that ensures ongoing communication between sub-cortical and cortical area along the entire process. The cyclic principle is also suitable to account for long-distance relations, movements and recursion.

## 5 Conclusion and further remarks

Based on the idea that language computational and representational processes are intrinsic to brain oscillatory activity, I developed a (preliminary) interdisciplinary formal proposal attempting to narrow the gap between linguistic and neuroscience for what concern the primitives assumed in syntactic computations (along the line of previous proposals: Murphy 2015a, 2016). This role assigned to oscillatory rhythms is justified by diverse body of evidence that signaling along reentrant cortical and cortico-thalamic areas paths is critical to cognition. Accordingly, within the model syntactic structures are derived by the specific functions assigned to cross-frequency coupled oscillatory rhythms: starting from the KS, the theta-gamma rhythms are cross-frequency coupled into the alpha-gamma-beta and into the gamma-beta-theta rhythms to generate the sentence along reentrant cortico-thalamic pathways through Merge, Label and Move operations. Crucially, this kind of model permits that both linguistic primitives and neurobiological primitives may be coherently investigated (an empirically tested) within a neurobiological perspective. The ultimate goal is to demonstrate whether cognitive brain functions are really represented by its oscillatory activity. This is probably the paradigm change that Mountcastle (1998) had announced for brain sciences toward the end of the last century, *pace* Chomsky (2000).

This perspective offers the basis to further reflect on the issue concerning what is special about language and its evolutionary genesis. Hauser, Chomsky & Fitch (2002: 1573) suggest that the Narrow Language Faculty – the computational mechanism of recursion – is recently evolved and unique to our species (recursion referring to a procedure that calls itself, or to a constituent that contains a constituent of the same kind). They propose that Narrow Language Faculty comprises only the core computational mechanisms of recursion as they appear in narrow syntax and the mappings to the interfaces with conceptual knowledge (and intentions) and perception-production mechanisms (see Pinker & Jackendoff 2005 for a critical discussion). More precisely, what the authors suggested is that a significant piece of the linguistic machinery entails recursive operations (Merge at least), and that these recursive operations must interface with SM and CI (and thus include aspects of phonology, formal semantics and the lexicon insofar as they satisfy the uniqueness condition of Narrow Language Faculty). Thus, the hypothesis focuses on a known property of human language that provides its most powerful and unusual signature: discrete infinity (Fitch, Hauser & Chomsky 2005: 182). The question is: how this property emerged? Is it an adaptive phenomenon – shared with other species and underwent refunctionalization at a certain evolutionary stage – or is unique to human language faculty and emerged by evolutionary selection? According to Hauser, Chomsky & Fitcht (2002), the latter hypothesis seems the more plausible.

As I have above discussed (cf. Section 3), reentrant neuronal activity leads to a kind of neuronal recursion: this represents a fundamental feature of thalamo-cortical activity characterizing only vertebrate nervous system (that is humans and other species). Reentrant activity represents not simple feedbacks but functions in a network as recursive multiple pathways, which update iteratively and hierarchically on a time scale of tens to hundreds of milliseconds, rapidly converging to the dynamic core’s synaptically connected neuronal network. It implies that computations in the brain assume primarily the form of interaction (if we accept early theoretical ideas that computation is interaction: Feynman 1996): i.e., intracellular interactions that generate recursive and integrated action potentials. Hence, it seems that recursive property is not exclusive to human brain and, most importantly, to human language.

If this primary mechanism is shared with non-human species, what is special to human language? What is absent in non-human vertebrate brains is the possibility to synchronize neuronal activity along a functional cortico-thalamic network: that is, for what concerns language computations, the synchronization of fronto-temporo-parietal cluster of neurons among themselves and with the thalamic nuclei (Edelman & Tononi, 2000). Reentrant activity per se is sufficient to generate primary conceptualization and categorization of the world (i.e., primary consciousness). However, high-order conceptualization and categorizations are possible only when long-term memory may be synchronously integrated with working memory to result in continuous computational and representational processes (i.e., secondary consciousness). This evolutionary specialization may be ascribed to functional synchronization and de-synchronization (coupling and uncoupling) of oscillatory rhythms that recursively bind together different conceptual objects (the words and all the relevant information) in a specie-specific recursive mechanism: that functional to build syntactic structures. So, some properties of the vertebrate brain may have been functionally reused during the emerging of unique cognitive abilities related to specific properties of natural languages: this may represent the Narrow Language Faculty recently evolved. While the hierarchy of brain rhythms themselves may be preserved, it is crucially their cross-frequency coupling relations which are at the basis of human language specialization (Murphy 2016b). I cannot address here the question whether Merge, Label or Concatenation plus Label (Hornstein 2009) represent the computational core of syntax computation, for which see Murphy (2015a,b; 2016a,b). But from the perspective outlined it seems problematic to exclusively assign to Merge the single computation operation, unique to language, to distinguishing it from other cognitive domains

To conclude, it is probable that synchronization of oscillatory rhythms has contributed to the emersion of a new connectivity that interconnected specific brain regions forming a fronto-temporo-parietal circuit that provides a more complete mechanism for richer representational capacities, viz. recursive capacities (Boeckx in press), together, of course, with the thalamo-basal ganglia loop.

## Acknowledgments

I thank Andrea Calabrese, Paolo Lorusso, Alec Marantz, Elliot Murphy, Leonardo Savoia for their comments on previous versions of the manuscript.

